# Rational tuning of a membrane-perforating antimicrobial peptide to selectively target membranes of different lipid composition

**DOI:** 10.1101/2020.11.01.364091

**Authors:** Charles H. Chen, Charles G. Starr, Shantanu Guha, William C. Wimley, Martin B. Ulmschneider, Jakob P. Ulmschneider

## Abstract

The use of designed antimicrobial peptides as drugs has been impeded by the absence of simple sequence-structure-function relationships and design rules. The likely cause is that many of these peptides permeabilize membranes via highly disordered, heterogeneous mechanisms, forming aggregates without well-defined tertiary or secondary structure. We demonstrate that the combination of high-throughput library screening with atomistic computer simulations can successfully address this challenge by tuning a previously developed general pore forming peptide into a selective pore former for different lipid types. A library of 2,916 peptides was designed based on the LDKA template. The library peptides were synthesized and screened using a high-throughput orthogonal vesicle leakage assay. Dyes of different sizes were entrapped inside vesicles with varying lipid composition to simultaneously screen for both pore size and affinity for negatively charged and neutral lipid membranes. From this screen, nine different LDKA variants that have unique activity were selected, sequenced, synthesized, and characterized. Despite the minor sequence changes, each of these peptides has unique functional properties, forming either small or large pores and being selective for either neutral or anionic lipid bilayers. Long-scale, unbiased atomistic molecular dynamics (MD) simulations directly reveal that rather than rigid, well-defined pores, these peptides can form a large repertoire of functional dynamic and heterogeneous aggregates, strongly affected by single mutations. Predicting the propensity to aggregate and assemble in a given environment from sequence alone holds the key to functional prediction of membrane permeabilization.

## Introduction

Recent years have seen a renewed interest in antimicrobial peptides (AMPs) as potential successors to small-molecule antibiotics^1-8^. The advantages of AMPs are clear: peptides are getting cheaper to synthesize on an industrial scale^9-13^, offer a near infinite chemical repertoire to target different species and cellular processes^14^, and can be rapidly screened using high-throughput methodologies^15-18^. AMPs are also proven, being a ubiquitous part of the innate immune defence of most branches of life. Although some AMPs are toxic to mammalian cells, many of these amphiphilic peptides selectively target and kill bacteria at low micro-molar concentrations without harming host cells^19-22^. Sequence analysis of >3,000 of known AMPs reveal a wide variation in amino acid composition, peptide length, and secondary structure; however, no clear functional motifs associated with antimicrobial activity have been identified to date, impeding rational optimisation and *de novo* design^2,23-25^. Despite this, there has been considerable progress in rational design and re-engineering of AMPs^24,26-35^. These studies have shown that a small number of amino acid mutations in a given sequence can significantly change functional properties such as pore stability^36^, antimicrobial activity^26,37-39^, pore size^30,36^, membrane selectivity^37^, and pH-dependent activity^27,40-42^. Peptide length also acts as an important factor. Ulrich *et al*. reported several rationally designed helical peptides with repeated KIAGKIA motifs with peptide length between 14 and 28 amino acids, and showed that the peptide length can affect its ability to penetrate and disrupt cell membranes^43,44^.

The ultimate goal is to develop AMPs that can selectively target specific membrane types in order to target pathogens with high potency, without harming host cells. A rational joint *in-silico*/experimental process has great potential for such *de novo* AMP design^45^. In the absence of reliable predictive rules for engineering the activity of membrane permeabilizing peptides, a recent breakthrough has been the use of synthetic molecular evolution, which is accomplished with orthogonal screening of a designed, iterative, combinatorial peptide library.^26,30,36,41,46^ Another strategy has been a simulation-guided design approach^24^, which we have applied to develop a potent pore-forming AMP starting from a membrane spanning polyleucine helix^47^. This new synthetic 14-residue AMP (sequence = GLLDLLKLLLKAAG), called LDKA, consists of only five amino acids (glycine, aspartic acid, lysine, leucine, and alanine) and shares similar sequences to many short antimicrobial peptides^24,48^. LDKA exhibits low micromolar antimicrobial activity and forms pores in both anionic POPG (1-palmitoyl-2-oleoyl-sn-glycero-3-phosphoglycerol) and neutral POPC (1-palmitoyl-2-oleoyl-sn-glycero-3-phosphocholine) lipid vesicles at low peptide-to-lipid ratios (P:L = 1:1000)^24^, which is comparable to the potent pore-forming peptide, melittin, and its gain-of-function analogue MelP5^36,37^.

Here, we explore whether we can rationally develop a general pore forming peptide into a selective pore former via a joint molecular dynamics (MD) simulation and an experimental library-screening approach. We start by tuning the hydrophobic moment and charge distribution to introduce preferential binding and pore formation in charged and neutral lipid bilayers and that this preferential binding correlates to activity against human versus bacterial cells. We demonstrate that relatively conservative sequence changes of the LDKA template can indeed modulate the induced preferential pore-forming potency in anionic versus neutral lipid bilayers as well as the size of the pores formed. We further demonstrate that these properties correlate well with antimicrobial activity for specific bacteria and selectivity for bacterial over human red blood cells.

Our results suggest that *in-vitro* activity, lipid selectivity, and aggregation propensities of AMPs depend highly on even the most conservative sequence changes. While the broad underlying properties correlate with simple descriptors that can be directly derived from the peptide sequence (e.g. hydrophobic moment, overall charge, and amphiphilicity), these quantities do not allow us to directly determine which sequence will be selective, or porate membranes at all. The peptides form a large repertoire of functional dynamic and heterogeneous structures in the membrane, and each sequence change can dramatically affect the oligomerization propensity, structure of the aggregates, ability to porate, and selectivity for different membrane compositions so desired for pharmaceutical application. This suggests that ultimately only structure (rather than sequence) based approaches, such as direct pore aggregation and equilibrium simulations, will enable predictive, rather than descriptive *de novo* AMP design.

## Results

### Rational peptide design

LDKA is a small pore-former in neutral POPC and anionic POPG vesicles and has low micromolar antimicrobial activity against bacteria. The goal of this library is to explore whether simple rearrangements of the LDKA sequence using four amino acids (Leu, Asp, Lys, and Ala), will allow modulation of pore-forming potential, pore-size, and targeting of specific membrane compositions. To achieve this, we have designed a combinatorial peptide library containing 2,916 LDKA analogues (**Figure 1a-b**). The LDKA template sequence was mutated in order to: (i.) adjust peptide hydrophobicity, (ii.) promote more salt bridge formation between the peptides, (iii.) introduce a central proline kink to the structure, and (iv.) substitute more positively charged residues on the C-terminus, which is one of the common motifs in the *Antimicrobial Peptide Database* (*APD;* http://aps.unmc.edu/AP)^49^.

**Figure 1.**
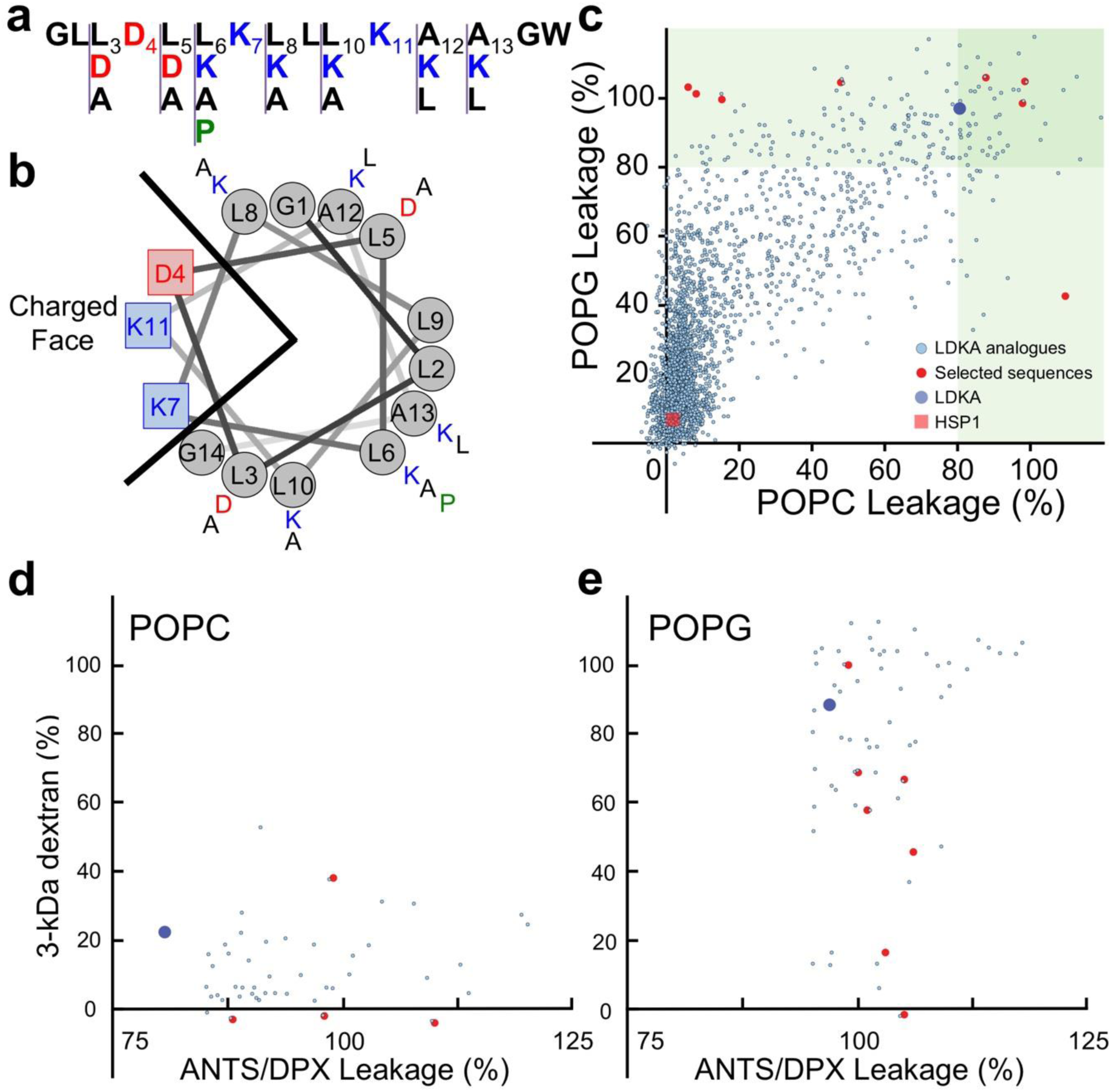
Peptide design and screening of the LDKA library contains 2,916 variants. **a**, Amino acid sequence of LDKA and its variants in the combinatorial peptide library. **b**, Helical wheel projection of LDKA showing charged and hydrophobic faces of the helix, which assumes it is a 100% helical configuration. Red and blue symbols present charged residues: negative charged and positive charged, respectively. Proline acts as a kink in the helix, and it is shown as green symbols. Other hydrophobic (leucine) and small (glycine and alanine) residues are indicated as grey symbols. **c**, High-throughput screen of LDKA peptide library induces fluorescent dye (ANTS/DPX) leakage from each POPC (x-axis) and POPG (y-axis) LUVs in 10 mM phosphate buffer at pH 7.0. Fluorescent dye release above 90% from POPC and POPG LUVs are highlighted in green areas, respectively, and the selected peptides were further analyzed their pore sizes using macromolecular fluorescent dye (3-kDa dextran). The selected LDKA library variants induce 3-kDa dextran releasing from each **d**, POPC and **e**, POPG LUVs.

Peptide hydrophobicity is modulated by interchanging leucine and alanine residues as well as substituting more positive (lysine) and negative charges (aspartic acid) in the sequence. The goal of these mutations is to fine-tune the peptide solubility and membrane-partitioning. To further allow for more structural plasticity of the peptide, we introduced a proline near the center of the peptide sequence, which is common in naturally occurring AMPs^50,51^. More charged residues (aspartic acid and lysine) were introduced to both facilitate inter-peptide salt-bridge formation and strengthen the peptide-peptide interface^52^, as well as to allow for a more polar central pore enabling larger multimeric channel structures^24,53^. Additional positive charges were introduced at the C-terminus to enhance peptide binding to anionic lipids, which is a common motif in many antimicrobial peptides from natural sources, such as Hylaseptin-P1^22,54^, Hylain 2^55^, melittin^56,57^, and maculatin^50,51^.

### Membrane specific poration and pore size

The potency and membrane selectivity of the 2,916 LDKA library peptides for zwitterionic (POPC) and anionic (POPG) large unilamellar vesicles (LUVs) was evaluated using a high-throughput liposome leakage screen. This approach allows us to detect and quantify the release of small fluorescent dye ANTS (8-aminonaphthalene-1,3,6-trisulfonic acid disodium salt; MW = 427 Da) and its fluorescent quencher DPX (p-xylene-bis-pyridinium bromide; MW = 422 Da) encapsulated in LUVs after addition of the library peptides. Neutral POPC LUVs serve as a simple model for mammalian membranes, while charged POPG LUVs serve as a very simplistic model for bacterial membranes enriched in anionic lipids.

**Figure 1c** demonstrates the fluorescent dye leakage fraction from both neutral and charged LUVs after addition of the library peptides. In this study, 11.2% of the LDKA analogues have POPG-favourable selectivity and induce >50% encapsulated dye leakage from charged POPG LUVs at low peptide concentration (P:L = 1:1000), while 0.4% cause leakage from neutral POPC LUVs only, and 6.6% disrupt both POPC and POPG LUVs. LDKA analogues that induce >90% dye leakage from POPC and POPG LUVs were screened for their ability to induce leakage of a larger 3-kDa TAMRA-biotin-dextran (TBD) dye^36^. Several LDKA-like peptides form larger pores in POPG vesicles, while the pores induced in POPC vesicles are generally smaller (**Figure 1d-e**).

Eight LDKA peptides with different lipid selectivity and pore sizes were selected from the high-throughput screen and sequenced using Edman degradation^58^. Table 1 shows that these peptides have 1 to 4 mutations compared to the LDKA template sequence. The most common mutation is leucine to alanine, occurring 13 times and in a total of 7 of the 8 peptides. Alanine to leucine occurred 6 times in 5 peptides, leucine to aspartic acid occurred 3 times in 3 peptides, and leucine to proline occurred once.

**Table 1.**
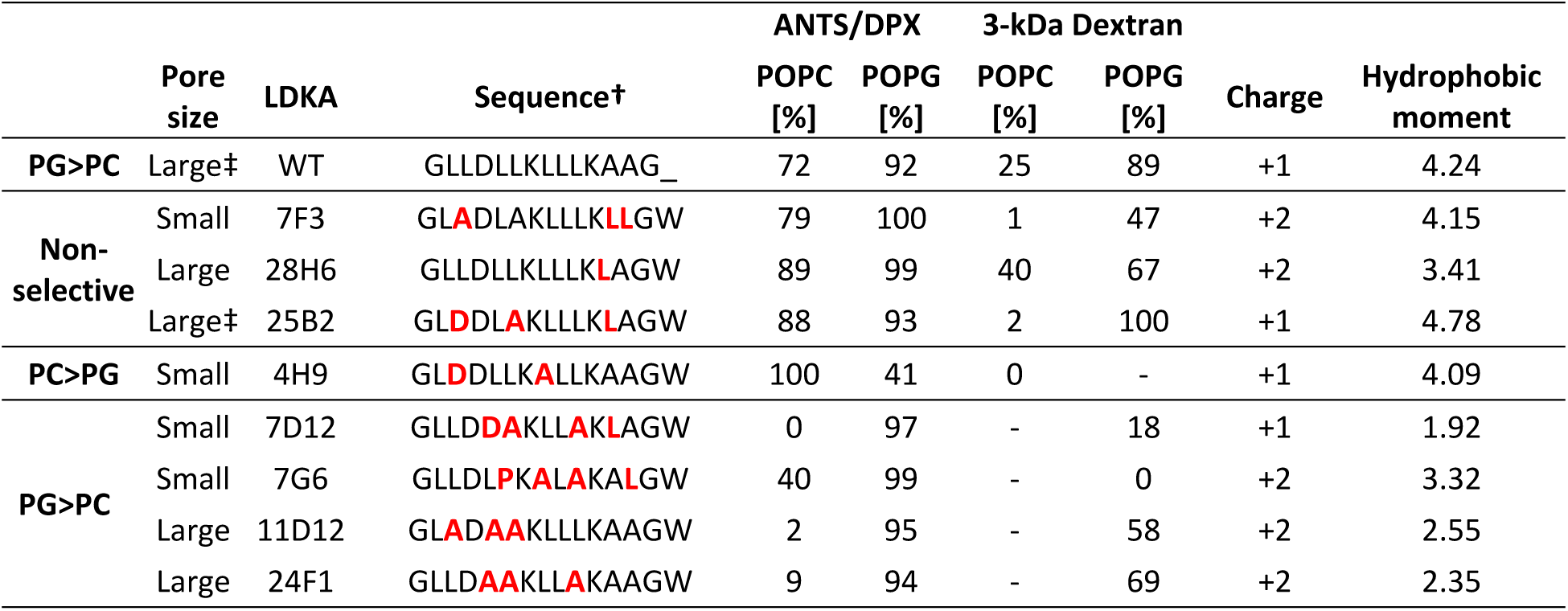
LDKA and its selected variants induce fluorescent dyes (ANTS/DPX and 3-kDa) release from each POPC and POPG LUVs with P:L = 1:1000 at pH 7 phosphate buffer. The fluorescent dye leakage fraction has been normalized to fit between 0 and 100 % by the positive control (LUV with potent peptide) and negative control (LUV with non-active peptide). †N-terminus is free, C-terminus: -NH_2_. ‡Large pore for PG only.

The analysis of selected peptide sequences showed positive-charged lysine is not a favourable substitution in the non-polar face of the LDKA template helix and the C-terminal motif (positions 6, 8, 10, 12, and 13). Instead, hydrophobic leucine and alanine are more preferable. This is in agreement with the evolutionary derivatives of 26-residue melittin, in that the positively-charged amino acids (lysine and arginine) are less likely to be favored in the non-polar face^26,37^.

Other than fixed lysine at positions 7 and 11, no additional lysine residues were observed in the analogues. Additional aspartic acids were observed at position 3 and 5, which is right next to the fixed aspartic acid at position 4 that can further promote salt bridge formation in peptide-peptide interactions. The net charges of these analogues are between +1 and +2, and they are consistent with the majority (net charge +1) of AMPs in the *APD*^23,49^. This shows that cationic residues can promote peptide binding to anionic bacterial membranes; however, more cationic charges may result in lower hydrophobicity and higher energy barriers to cross the hydrophobic core of membranes. Therefore, a longer peptide length is needed to strengthen the hydrophobicity when the sequence contains more charges. A natural membrane-active peptide, melittin (sequence: GIGAVLKVLTTGLPALISWIKRKRQQ-Amide), is a good example. Although it has four positive charges (-KRKR-) in its C-terminus, longer peptide length (26 amino acids) and the hydrophobic N-terminus (GIGAVLKVL-) make it hydrophobic enough to span cell membranes.

Table 1 reveals that leucine to alanine mutations are generally sufficient to prevent poration in neutral POPC membranes, while the peptides still porate charged POPG membranes, which is similar to the L16G mutation of melittin^37^. More specifically, the LDKA analogues that only induce ANTS/DPX leakage from anionic POPG LUVs have 4-5 leucines, while the analogues that can porate both POPC and POPG LUVs have 6-7 leucines in their sequences. The net charge of all LDKA wildtype and analogues are between +1 and +2, and we did not observe any anionic peptide, neutral peptide, or peptide that has net charge greater than +2. This suggests that the membrane-selectivity is driven by hydrophobic moment to POPC but including electrostatics on POPG.

### Binding to mixed membranes

To investigate the root cause of the different leakage preferences of LDKA analogues for POPC and POPG membranes, we studied the binding and secondary structural properties of LDKA analogues using tryptophan fluorescence and circular dichroism (CD) spectroscopy, respectively. Peptide solutions (50 µM peptide concentration) were titrated with POPC and POPG LUVs (between 0-5 mM) and the corresponding changes of tryptophan fluorescent spectra were collected, yielding binding free energies and helicities of the peptides, albeit without any structural information on the underlying poration process (**Figure S1**, supplement).

Further studies were performed to answer why some peptides (i.e. 7D12, 7G6, 28H6, 11D12, and 24F1) show selectivity for either membrane type. First, we characterised peptide secondary structural changes and binding to LUVs containing binary mixtures of POPC and POPG lipids. **Figure 2** (7D12, 7G6, and 28H6) and **Figure S5** (11D12 and 24F1) show changes in the tryptophan fluorescence and CD spectra for these peptides upon addition of LUVs for whom the ratio of POPG was elevated from 0 to 100% with 20% increments (0, 20, 40, 60, 80, and 100% POPG). These analogues are sensitive to the anionic POPG lipid and have significant structural change with small PG fraction (20% POPG), except 7D12 which is less sensitive to anionic lipid. These membrane selective peptides only bind to POPG and show little or no binding to POPC, which is consistent to our liposome leakage assay.

**Figure 2.**
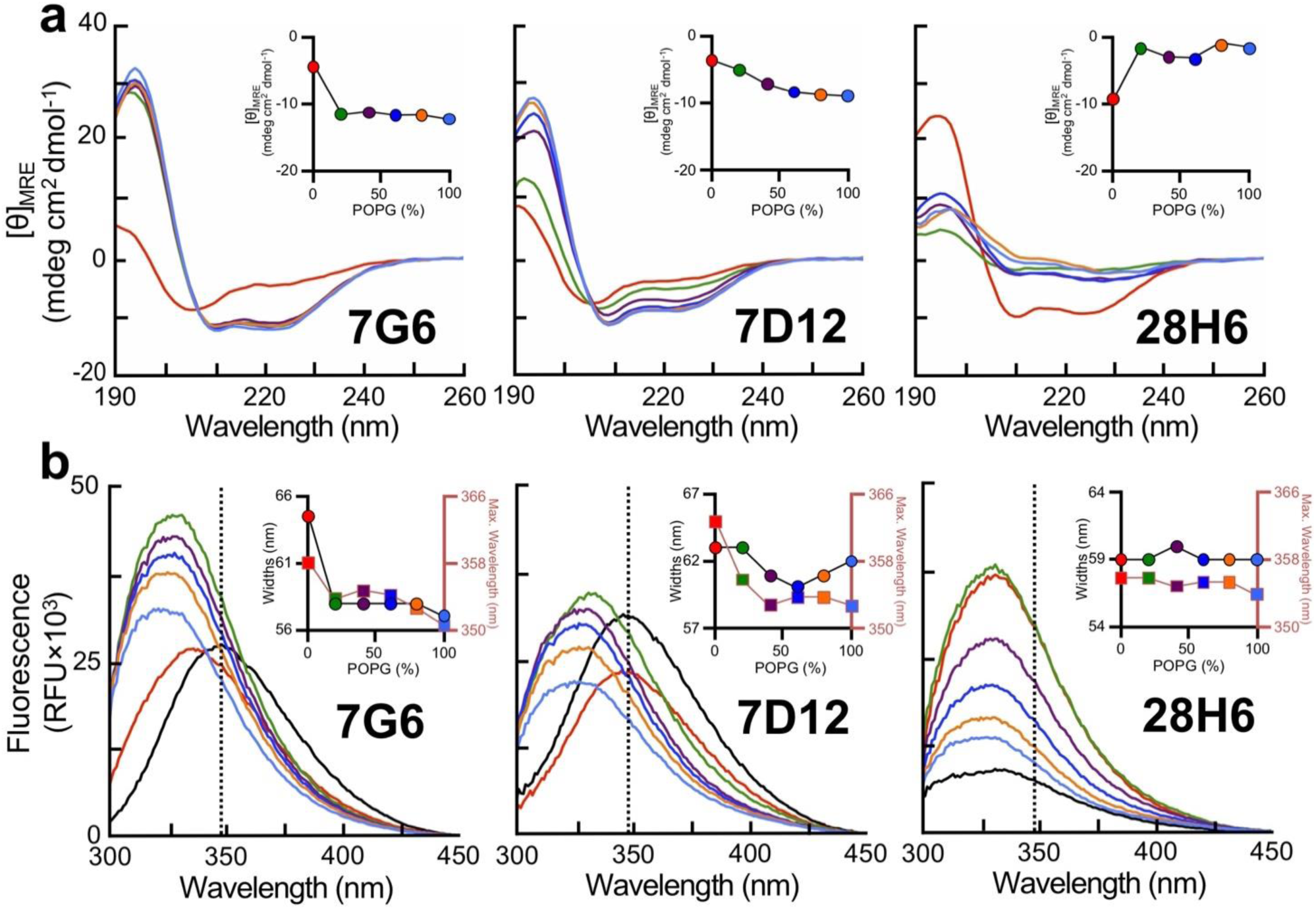
Peptide binding and folding of 7G6 (left), 7D12 (middle), and 28H6 (right) onto binary mixtures of charged lipid (POPG) and neutral lipid (POPC) LUVs. **a**, Circular dichroism spectroscopy and **b**, tryptophan fluorescent binding assay of LDKA peptides (50 µM) at P:L = 1:12 in 600 µM POPC/POPG LUVs with different lipid compositions: no lipid (black), 100% POPC (red), 80% POPC and 20% POPG (green), 60% POPC and 40% POPG (purple), 40% POPC and 60% POPG (dark blue), 20% POPC and 80% POPG (orange), and 100% POPG (light blue). The experiments were performed in 10 mM phosphate buffer at pH 7.0.

### Simulations of two similar sequences

The information gained from the experimental screen is limited in that there is an absence of a nuanced correlation between simple peptide descriptors and selectivity and leakage propensity. Since a multitude of AMP structures can cause membrane permeabilization,^14^ it is critical to identify which overall mechanism applies for the chosen library template. Without knowledge of pore structures in the membrane, it is difficult to explain the role of individual sequence changes on both selectivity and poration ability, rendering the design process blind. Here, computer simulations offer to fill in the missing information. We have demonstrated before for numerous peptide/membrane systems that long-scale equilibrium MD simulations are now able to directly generate aggregate structures in different membrane types from peptide sequence alone^45. 53^ The computational effort is – for now – enormous, so only a small subset of the LDKA library was chosen for structural investigation, focusing on 2 peptides that are almost similar but have very different membrane selectivity: 25B2 (sequence: GLDDLAKLLLKLAGW-Amide) and 7D12 (sequence: GLLDDAKLLAKLAGW-Amide). Despite very similar sequences, 25B2 (toxic peptide) causes fluorescent dye leakage from both POPC and POPG LUVs, and 7D12 (membrane-selective peptide) only porates POPG LUVs without disturbing POPC membranes. We sought to observe how these library peptides lyse their target membranes, and how almost identical sequences can have vastly different binding properties.

Both peptides have a net charge +1 and have the same C-terminal motif (-KLAGW-Amide). The only differences are (i.) aspartic acid shifts from position 3 in 25B2 to position 5 in 7D12, and (ii.) hydrophobic position 10 where 25B2 is leucine and 7D12 is alanine. A quick analysis shows these simple modifications result in a hydrophobic dipole moment of 4.8 in 25B2 and 1.9 in 7D12. Mirroring the biophysical experiments, we performed peptide-assembly simulations of 7D12 and 25B2 in both POPC and POPG bilayers. (**Figure 3 and Table S3**). Simulations and experiments show that 25B2 results in higher helical content than 7D12 (**Table S2 and Table S3**). Similar to our prior simulations of LDKA, the peptides spontaneously insert and form a large number of heterogeneous oligomeric pore-structures. These can range from 3-9 peptides, with a core of mainly 4-5 tilted TM inserted peptides, supported by several surface-bound peptides that are more loosely attached. Since the sequences are short, the main arrangements are strongly tilted and double-stacked, rather than a membrane-spanning barrel-stave layout. Peptides align both parallel and anti-parallel at various levels of insertion. The large number of charged sidechains, both cationic and anionic, enable small water-filled bilayer channels with many cross-peptide salt-bridges, pulling in both lipid headgroups and ions. Peptides usually leave and join these small aggregates, resulting in no overall stable structures but rather in a wide variety of different pore assemblies. There is substantial water and ion flux across these, with higher oligomers yielding larger flux. Both cations and anions can translocate across the pores, with a preference for cations in the POPG simulations, presumably due to the more anionic environment of the pore aggregates, where PG headgroups are pulled into the membrane. The heterogenous nature of the pore aggregates indicates a highly dynamic equilibrium which is strongly influenced by individual sequence changes. 7D12 is shown to be selective: It does not insert and form aggregates in POPC, but remains on the surface, indicating that pore aggregates are not stable in this membrane, and the surface state is preferred.

**Figure 3.**
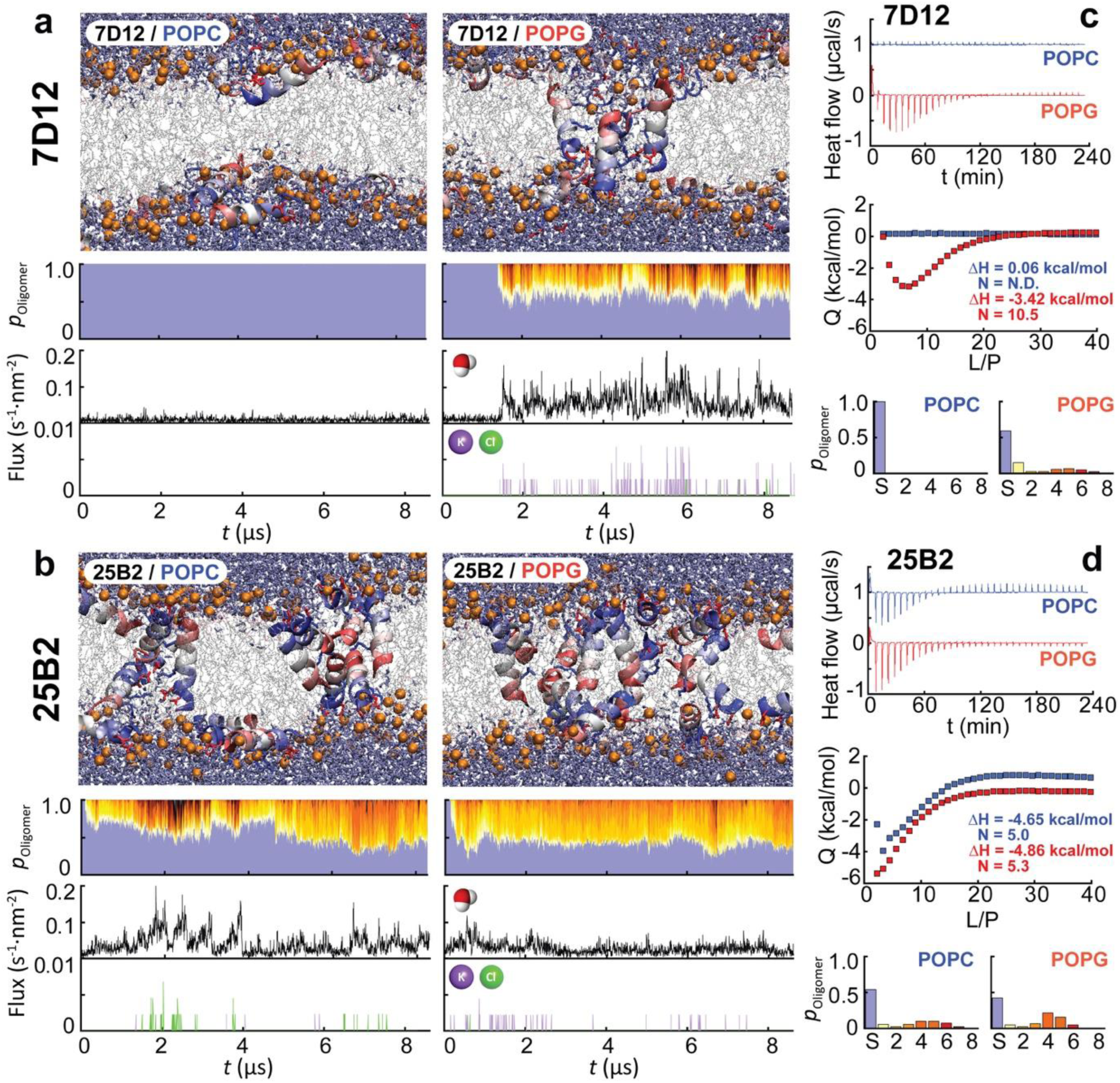
Multi-microsecond molecular dynamics simulations reveal the spontaneous self-assembly of the membrane-selective peptide 7D12 and the toxic peptide 25B2 and their oligomeric structural ensembles in POPC and POPG membranes. **a** and **b**, Representative pore aggregates (peptides colored blue (N-) to red (C-) terminal, lipid phosphates as orange beads), oligomeric occupation plots (blue = S-state, yellow = single TM, red-dark = higher TM oligomers, overall distribution on right), and cross-membrane water and ion flux caused by the pore assemblies. **c** and **d**, Isothermal titration calorimetry of the heat release/absorption of the peptide-lipid interactions. The integrated ITC data curve of 7D12 and 25B2 with each POPC and POPG LUVs is also shown. The concentration is fixed at 100 µM with titrated lipid LUVs in 10 mM phosphate buffer at pH 7.0. The ITC data is consistent with the simulation results for the binding selectivity of 7D12 for POPG.

### Isothermal titration calorimetry

To directly compare the above simulations, we applied isothermal titration calorimetry (ITC) to further characterize their thermodynamic parameters: enthalpy (*∆H*) and stoichiometry (*N*). Titration of POPC vesicle into membrane-selective peptide 7D12 (100 µM peptide concentration) results in *∆H* = 0.1 kcal/mol (**Figure 3c-d**), which is consistent with the tryptophan fluorescent binding assay that it is does not bind strongly to POPC vesicle (**Figure S1**). Titrating POPG vesicle into 100 µM 7D12 has significant heat release (*∆H* = −3.4 kcal/mol) with the stoichiometry of 11 lipids per peptide (*N* = 11). On the other hand, the toxic peptide 25B2 with titrated POPC and POPG vesicles shows *∆H* = -(4.7-4.9) kcal/mol, and they both have the same stoichiometry of *N* = 5 lipids per peptide.

The results of MD simulations and ITC are consistent. 7D12 in POPG, and 25B2 in both membrane types assembled channel-like architectures in MD simulations and showed significant heat release in ITC. In contrast, 7D12 in POPC bilayers neither formed any structure, nor induced any heat release/absorption. Thus, there is a remarkable agreement between experiments and simulations. The lower hydrophobic moment of 7D12 appears to explain the less thermostable helical structures than other peptides (**Figure S2**), and the unfolded structures are more disordered than the helical structure of 25B2 as compared to what we observed in ITC (**Figure 3c-d**). Therefore, it suggests hydrophobic moment is a determinant to promote membrane selectivity.

### Hemolysis and antibacterial activity

To test toxicity of the LDKA analogues against human cells, we performed a hemolysis assay (**Figure 4a**). LDKA wildtype is hemolytic at moderate micromolar concentrations with a hemolytic concentration lysing 50% of red blood cells (HC_50_) of 55.1 µM (Table 2). The peptide-induced POPC LUV leakage is correlated with the hemolytic activity (**Figure 4b-c**). The peptides that induce leakage from POPC LUV at low peptide concentration (P:L = 1:1000) are hemolytic (HC_50_ = 1-57 µM). More specifically, 7F3 (HC_50_ = 1.1 µM) and 28H6 (HC_50_ =1.2 µM) are as powerful as natural toxin melittin and its gain-of-function derivative MelP5 (HC_50_ = 1-3 µM)^37^. All POPG-favourable peptides have no effect to human red blood cell, even at 75 µM peptide concentration.

**Table 2.**
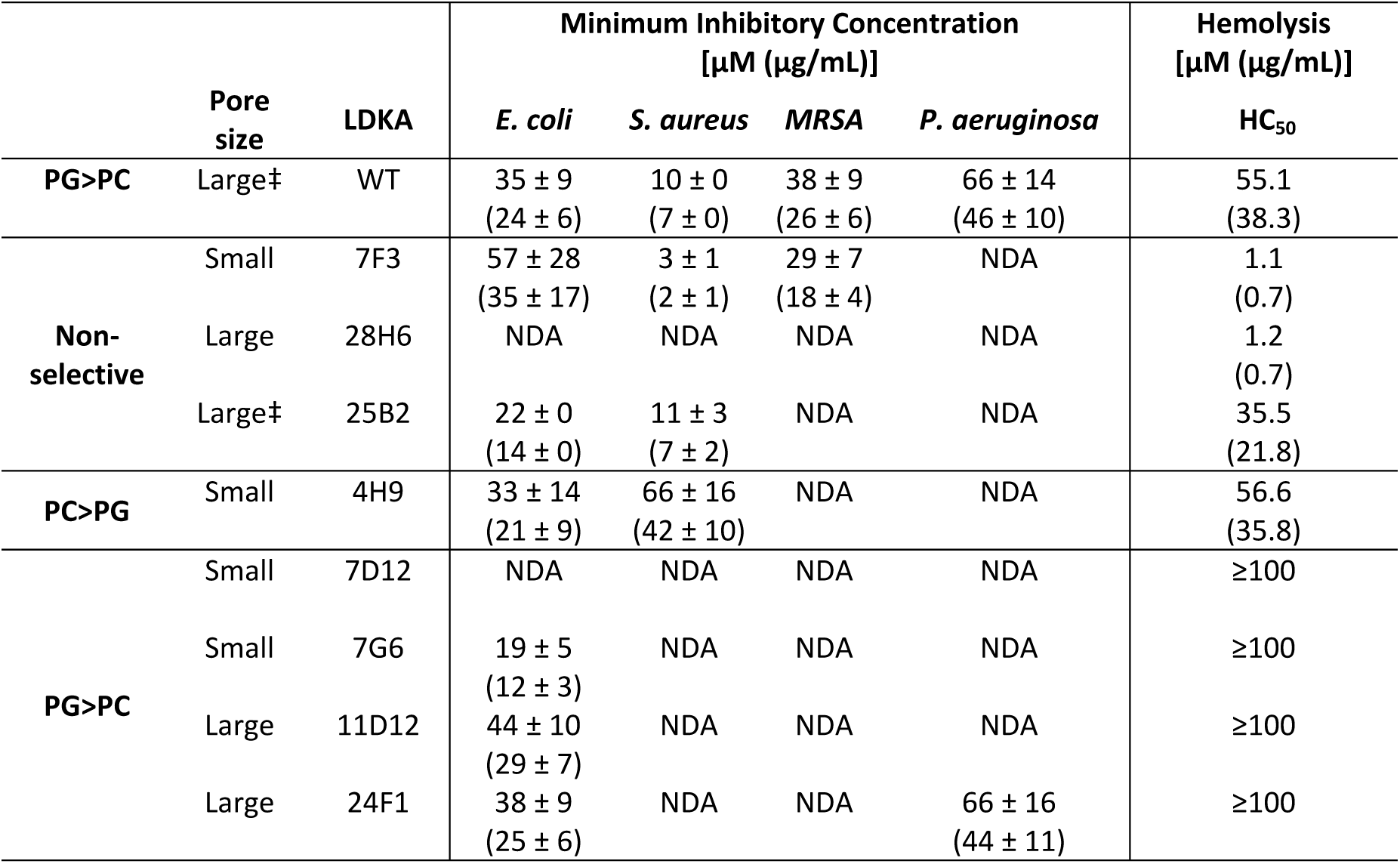
*In-vitro* experiments of LDKA analogues show their minimum inhibitory concentration with *E. coli, S. aureus*, and *P. aeruginosa*, and hemolysis shows their hemolytic activity at the corresponding peptide concentrations. HC_50_ present the hemolytic activity of peptide concentration to kill 50% of human red blood cell. 75 µM peptide concentration is the maximum amount that were tested. “NDA” means “not determinable”. ‡Large pore for PG only.

**Figure 4.**
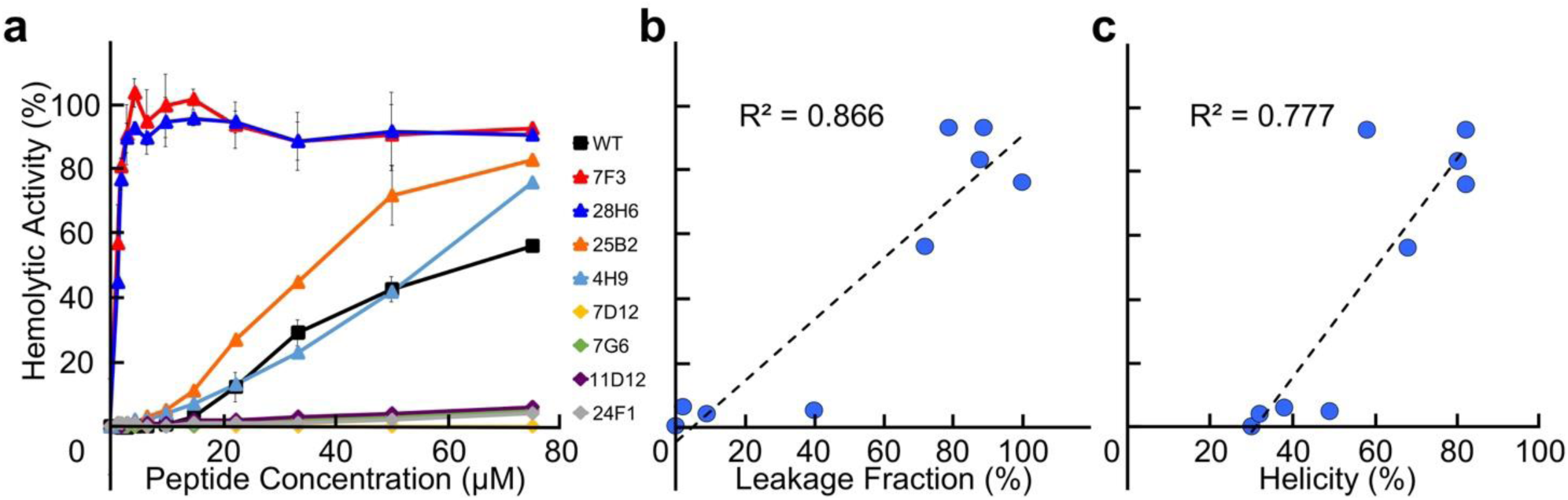
Selected LDKA analogues and their *in vitro* hemolytic activity with human red blood cell. **a**, Hemolytic activity with human red blood cell varies in peptide concentration. Linear regression analysis of **b**, hemolysis (at 75 µM peptide concentration) versus ANTS/DPX leakage fraction from POPC LUV (at P:L = 1:1000). y = 0.954x - 4.540 and R^2^ = 0.866, where x = ANTS/DPX leakage fraction (%) and y = hemolytic activity (%). **c**, Linear regression analysis of hemolytic fraction (at 75 µM peptide concentration) versus peptide helicity in POPC LUV (at P:L = 1:12). y = 1.717x - 52.791; R^2^ = 0.777, where x = helicity (%) and y = hemolytic activity (%).

The real test is how selectively the selected peptides target and kill various bacteria. The antibacterial activity of LDKA analogues against *E. coli, S. aureus*, and *P. aeruginosa* was tested *in vitro* in nutritionally rich medium. LDKA wildtype inhibits growth of all three bacteria at micromolar peptide concentrations of a similar range to potent AMPs from natural sources. From our screen, most of the POPG-favourable peptides (7G6, 11D12, and 24F1) have antibacterial activity and specificity against *E. coli* with 19-44 µM minimum inhibitory concentration (MIC) but are not active against other bacterial species: *S. aureus* and *P. aeruginosa* (Table 2). The toxic peptides 7F3, 25B2, and 4H9 are effective inhibitors against *E. coli* and *S. aureus*, but not *P. aeruginosa*. The results show that these peptides have specificity toward different bacterial species.

### Activity against antibiotic-resistant strains

Bacteria can mutate and develop resistance against conventional antibiotics^2,59-61^, which mostly target specific proteins, ribosomes, or DNA. Antimicrobial peptides exert their effects through a more generalized mechanism: membrane poration. We selected four conventional antibiotics that have different mechanisms to kill bacteria: ceftazidime, ciprofloxacin, streptomycin, and gentamicin. Ceftazidime interferes with bacterial cell wall formation^62,63^. Ciprofloxacin inhibits DNA gyrase, type II topoisomerase, and topoisomerase IV to separate bacterial DNA and DNA replication, thus inhibiting cell division^64^. Streptomycin and gentamicin inhibit protein synthesis at the ribosome^65,66^. Drug-resistant *E. coli* strain ATCC 25922 cultures were grown in the presence of each antibiotic at elevated concentration and the surviving strains was selected to grow for 10 generations, resulting in a 4 to 16-fold resistance to these antibiotics compared to their 1^st^ generation strain (**Figure 5a**).

**Figure 5.**
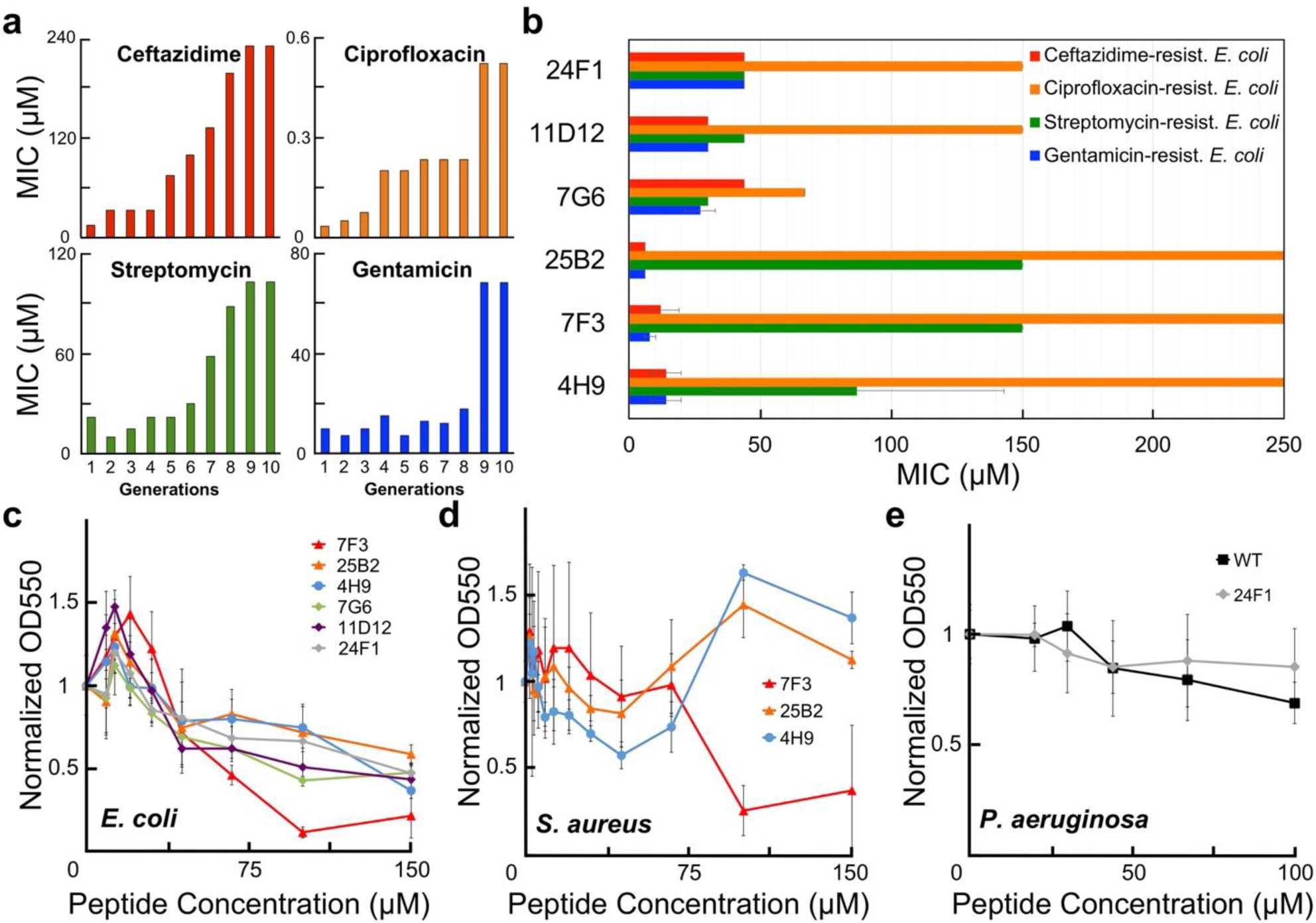
LDKA analogues against four different drug resistant *E. coli* strains and bacterial biofilms. **a**, Minimum inhibitory concentrations (MICs) of four conventional antibiotics (ceftazidime, ciprofloxacin, streptomycin, and gentamicin) were treated with serial *E. coli* generations. The *E. coli* that survives below/near the MICs was selected for the next generation. **b**, MICs of LDKA analogues (membrane-selective peptides: 24F1, 11D12 and 7G6; toxin peptides: 25B2, 7F3 and 4H9) against four different strains of drug resistant *E. coli*. Antibacterial activity of LDKA analogues against quantitative biofilm formation on polystyrene 96-well plate for 3 hr treatment. Selected analogues were tested with each **c**, *Escherichia coli* biofilm, **d**, *Staphylococcus aureus* biofilm, and **e**, *Pseudomonas aeruginosa* biofilm.

LDKA analogues were tested against these four drug-resistant *E. coli* cultures. Membrane-selective analogues (7G6, 11D12, and 24F1) remain effective and consistently inhibit the growth of ceftazidime-resistant, streptomycin-resistant, and gentamicin-resistant *E. coli* strains with 27-44 µM peptide concentrations (**Figure 5b**). Toxic peptides (4H9, 7F3, and 25B2) are effective against ceftazidime-resistant and gentamicin-resistant *E. coli* at low peptide concentrations (6-14 µM). Surprisingly, none of the peptides are effective against the ciprofloxacin-resistant *E. coli* strain.

### Activity against biofilms

In clinical settings, bacteria are mostly found in biofilms that are the key drivers of infections^67,68^. We therefore challenged our LDKA analogues against bacterial biofilms, which are generally much more resistant than planktonic equivalents^69^. The results showed that the selected LDKA analogues (4H9, 7F3, 25B2, 7G6, 11D12, and 24F1) can eliminate ∼50% of the *E. coli* biofilm in the presence of 67-150 µM peptide. Only 7F3 is capable of reducing *S. aureus* biofilms by ∼50% with 100 µM peptide concentration, and none of the analogues work against *P. aeruginosa* biofilms (**Figure 5c-e**).

## Discussion and Conclusions

In this study, we used the leucine-rich membrane-active peptide LDKA as a sequence template in order to test whether the combination of database-guided combinatorial peptide library screening, and direct MD simulation of membrane aggregation, can tune the template to significantly change its secondary structure, potency, and membrane specificity. Similar rational combinatorial design has been used before to develop and tune the activity of other AMPs^26,37,50,70^. The LDKA library peptides were designed using only four different amino acids (Asp, Lys, Leu, and Ala) in a template sequence of GxxDxxKxxxKxxGW-Amide, where ‘x’ represents one of the four amino acids. Our LDKA analogues reveal that a small number of conservative substitutions (Leu to Ala) in the LDKA sequence can dramatically change the selectivity toward different membrane types (anionic and neutral LUVs), resulting in specificity to different bacteria species and human red blood cells. This is consistent with Krauson *et al*., who showed a single-residue change of a leucine at position 16 to glycine (L16G) can redirect the general toxicity of melittin towards bacteria only, leaving red blood cells unharmed^37^. A similar study introduced charged amino acids in the C-terminal motif of HYL-20 peptide, fine-tuning the selectivity against several bacteria strains with negligible hemolytic activity^70^. The fact that we did not observe this feature in our LDKA peptide library suggests, again, that simple generic structure-function rules are not applicable to membrane active peptides.

The dependence of the drastic changes in selectivity and leakage propensity upon small sequence changes demonstrates the limitation of overall macroscopic peptide descriptors such as hydrophobic moment and polar angle as design criteria. For example, the hydrophobic moment is somewhat correlated to hemolytic activity (**Figure S4d**), and the LDKA peptide library suggests a hydrophobic moment of 3.37 as a cut-off for toxicity toward human red blood cells; however, this does not apply to 26-residue peptide melittin (hydrophobic moment = 3.94) and its membrane-selective analogue (hydrophobic moment = 3.44-3.46). Hydrophobic moment estimates could be limited as they are based on a single, perfectly helical peptides, and do not consider peptide-peptide aggregates and assemblies, as observed in our MD simulations.^71-76^ Experimentally, fluorescent dye leakage from POPC vesicle is also a reliable model to predict the hemolytic activity with a linear correlation (R-squared value = 0.87; **Figure 4b**). It is similar to structure-function relationship that shows R-squared value 0.78 between helicity in POPC LUV and hemolysis (**Figure 4c**).

The absence of strong correlations between macroscopic peptide descriptors and selectivity and leakage propensity means detailed structural models are needed to show what is going on. Advances in computer performance have enabled long-scale (multi-µs), fully atomistic MD simulations to provide that picture.^53,77-87^ We have demonstrated before that MD simulations are now able to directly predict aggregate structures in membranes.^45,53,87^ The simulations here show the atomic details of how these short membrane-spanning peptides selectively fold, aggregate, and form water pores in specific lipid bilayers (**Figure 3**). Key to these simulations is that they are not stuck in initial conditions. The pore aggregates are predicted without any prior information and fluctuate sufficiently to reveal the major structural assemblies that these peptides are expected to populate at either equilibrium, or during a membrane permeabilization event. Structures are highly heterogenous. The selectivity found in the experiments is reproduced in the simulations: 7D12 only folds and assembles in anionic POPG bilayers. There are no TM pores for 7D12 in POPC, with the peptides staying on the membrane surface and no noticeable water leakage. This is consistent to the experimental findings, and it demonstrates the extreme effect even tiny sequence changes can have on the pore forming equilibrium. Anionic sidechains are known for their steep insertion penalties,^88^ so the reason for the lack of TM insertion likely is the shift of the 2 Asp residues one position towards the center of the peptide. The propensity of a peptide sequence to aggregate and assemble in a given environment depends in a highly complex and non-linear way on the its sequence. Therefore, purely sequence-based design approaches are likely not suited for peptides that can form such a large repertoire of functional structures.

How do the designed library peptides perform in killing pathogens? Several minor mutations of LDKA can fine-tune the potency and specificity to kill *E. coli*, but not harming human red blood cells, *S. aureus*, and *P. aeruginosa*. Most of the LDKA peptides are able to inhibit the growth of *E. coli* with 19-57 µM peptide concentrations, except 28H6 and 7D12 that fail to treat *E. coli*. 7D12 has the lowest hydrophobic moment 1.92 that may not be strong enough to fold and assemble in the bacterial membrane. Furthermore, the surface protease OmpT on the outer membrane of the *E. coli* may confer resistance to these leucine-rich peptides by cleaving their peptide bonds and degrading the peptides^89^. Our study suggests that small fluorescent dye leakage assay with POPC LUVs and POPG LUVs are ideal models to quickly screen the AMPs for their hemolytic activity and antibacterial activity. However, formation of different pore size does not correlate to the *in-vitro* activity in our study.

Antibiotic-resistance is another serious threat. Half of the LDKA analogues that are able to inhibit the *S. aureus*, but many of them fail to eliminate the super bug, methicillin-resistant *Staphylococcus aureus* (MRSA) (Table 2). Although our LDKA analogues are less or not effective against *S. aureus* and *P. aeruginosa*, many of them are useful to eradicate *E. coli* with negligible effect to human red blood cell. Remarkably, these LDKA peptides are active against drug-resistant *E. coli* (**Figure 5a-b**) and biofilm (**Figure 5c-e**) with micromolar peptide concentrations.

Our study demonstrates a simple methodology of the rational design of membrane-selective peptides, revealing the potential of using MD simulations to fine-tune the membrane selectivity for peptide design and protein engineering for different cell types. This demonstrates the feasibility of computer-guided antibiotics design^24,90-92^, developing potent antimicrobial peptides that have effective membrane selectivity to distinguish between human red blood cells and bacterial membranes, and even between different bacterial species. The key advantage of *in-silico* techniques is the vastly larger combinatorial space that can be explored in comparison to experimental library screening. In this study, the large-scale all-atomistic simulation effort was limited to only a few sequences and target membranes due to the heavy resources required. However, the strong correlation to the experimental results demonstrates the maturity of these techniques. With rising computing power in the near future, the library screening effort will be shifted towards the computational side. This combined experimental/computational approach opens the path to apply these LDKA analogues, and numerous other designed peptides to various different biomedical applications, e.g. antibiotics, biosensors, and drug delivery.

## Methods

### Combinatorial peptide library synthesis

The synthesis of combinatorial peptide library was modified from the method described by Krauson, *et al*^26^. Peptides library synthesis was performed using Tentagel® NH_2_ macrobeads (280-320 µm bead diameter) particle size (∼65,550 beads/g) using Fmoc solid-phase peptide synthesis. Each bead only has one peptide sequence. A photolinker is attached between peptide and bead to allow the UV light-induced cleavage of homogenous peptide from bead in each well. The quality of the peptide library was verified by mass spectrometry (e.g. MALDI) and Edman sequencing. After placing one bead in each well of 96-well microplate, the photolinker between peptide and bead was cleaved with 5 hr of low-power UV light on dry bead, which were spreading to a dispersed single layer in a glass dish. The peptides were each dissolved in DMSO, quantified by tryptophan absorbance using nanodrop, and stored in −20 °C freezer.

### Bulk peptides and chemicals

The selected LDKA-like peptides were synthesized using standard Fmoc chemistry and purified to 98% purity using reverse phase HPLC by GenScript, Inc (Piscataway, NJ, USA). The N-terminus was positively charged amine group and C-terminus is neutral amide group. Peptide purity and identity were confirmed by HPLC and ESI mass spectrometry. The solubility test was performed by GenScript, Inc (Table S1).

### Large unilamellar vesicle (LUV) preparation

1-palmitoyl-2-oleoyl-sn-glycero-3-phosphocholine(POPC),1-palmitoyl-2-oleoyl-sn-glycero-3-phospho-(1’-rac-glycerol) (POPG), and hexadecanoyl sphingomyelin (Egg SM; PSM) were purchased from Avanti Polar Lipids. Lipids were dissolved in chloroform, mixed, and dried under nitrogen gas in a glass vial. Any remaining chloroform was removed under vacuum overnight. To make LUVs lipids were resuspended in 10 mM sodium phosphate buffer (pH = 7) with 100 mM potassium chloride. LUVs were generated by extruding the lipid suspension 10 times through 0.1 µm nuclepore polycarbonate filters to give LUVs of 100 nm diameter^93^.

### ANTS/DPX leakage assay

5 mM ANTS and 12.5 mM DPX were entrapped in 0.1 µm diameter extruded vesicles with lipids^94,95^. Gel filtration chromatography of Sephadex G-100 (GE Healthcare Life Sciences Inc) was used to remove external free ANTS/DPX from LUVs with entrapped contents. LUVs were diluted to 0.5 mM and used to measure the leakage activity by addition of aliquots of LDKA. Leakage was measured after 3 h incubation. 10% Triton was used as the positive control to measure the maximum leakage of the vesicle. Fluorescence emission spectra were recorded using excitation and emission wavelength of 350 nm and 510 nm for ANTS/DPX using a BioTek Synergy H1 Hybrid Multi-Mode Reader.

### Macromolecule release assay

Several different size dextrans were prepared and labelled with both TAMRA and biotin. Conjugated dextran was entrapped in POPC LUVs as described above^36^. External dextran was removed by incubation with immobilized streptavidin. Streptavidin labelled with an Alexa-488 fluorophore was added during the leakage experiment with the peptide. The sample was incubated for 3 hours before measuring Alexa-488 fluorescence. A control without added peptide served as the 0% leakage signal and addition of 0.05% vol. Triton X-100 was used to determine 100% leakage.

### Circular dichroism (CD) spectroscopy

LDKA solutions (50 μM) in 10 mM phosphate buffer (pH 7.0) were co-incubated with 800 μM POPC:POPG (1:1) and POPC:CHOL (7:3) LUVs in identical buffer (see *LUV preparation* above). CD spectra were recorded using the synchrotron radiation circular dichroism beamline on ASTRID at Aarhus University. Spectra were recorded from 270 to 170 nm with a step size of Δλ = 0.5 nm, a bandwidth of 0.5 nm, and a dwell time of 2 s. Each spectrum was averaged over 3 repeat scans. The averaged spectra were normalized to molar ellipticity per residue. The raw data were analyzed using DichroWeb <http://dichroweb.cryst.bbk.ac.uk/>.^96-98^

### Peptide thermostability enables advanced sampling at high temperatures

The LDKA peptide is resistant to thermal denaturation when bound to the membrane and the simulated helicity is comparable to the experiments (Table S2, Table S3, and **Figure S2**)^24,47,53,99-102^. This allows all simulation to be run at 120 °C, increasing pore-formation kinetics. We have previously demonstrated that elevating the temperature does not change conformational equilibria or partitioning free energies of helical membrane-active peptides, provided they are stable against thermal denaturation (see Supplement); however, the vast increase in sampling kinetics at high temperatures allows simulation of peptide folding, bilayer partitioning, and pore assembly without the need for advanced sampling techniques that require additional information or may bias the system^24,47,53,99-103^.

### Tryptophan fluorescent binding assay

The protocol was modified from the original method described by Christiaens, et al.^104^. LDKA peptides (50 µM) and POPC/POPG (600 µM) were prepared in 10 mM phosphate buffer (pH 7.0). The solutions were incubated and measured after 60 minutes. Excitation was fixed at 280 nm (slit 9 nm) and emission was collected from 300 to 450 nm (slit 9 nm). The spectra were recorded using Synergy H1 Hybrid Multi-Mode Reader and Cytation™ 5 Cell Imaging Multi-Mode Reader from BioTek and were averaged by 3 scans.

### Bacterial minimum inhibitory concentration (MIC)

*Escherichia coli* strain ATCC 25922, *Staphylococcus aureus* strain ATCC 25923, and *Pseudomonas aeruginosa* strain ATCC PAO1 were used in this study. Overnight cultures were sub-cultured and diluted to an initial bacterial cell density of ∼3 x 10^5^ colony forming units (CFU) per mL in Lysogeny broth. Cell counts were determined by measuring optical density at 600 nm (OD_600_), with an optimal sensitivity at OD_600_ = 0.3-0.6 in a 1 cm path-length cuvette. OD_600_ = 1 corresponds to 1.5 × 10^8^ CFU/mL for *S. aureus*, 5 × 10^8^ CFU/mL for *E. coli*, and 2.04 × 10^8^ CFU/mL for *P. aeruginosa*. Bacteria were added to peptide (LDKA and indolicidin) dilutions (1.3, 2.0, 2.9, 4.4, 6.6, 9.9, 14.8, 22.2, 33.3, 50.0, and 75.0 µM) and co-incubated at 37 °C. After 12 hr incubation, the optical density of the wells were recorded on a plate reader to determine whether they were sterilized (OD_600_ < 0.08) or were at stationary phase growth (OD_600_ > 0.5). Intermediate values, which were rare, were considered positive for growth. Average minimum sterilizing concentrations were calculated from the lowest peptide concentration that sterilized the bacteria in each serial dilution. The samples were done in sextuplet.

### Biofilm

The formation of biofilm and quantification was modified from the method described by O’Toole, et al.^105^. *Escherichia coli* strain ATCC 25922, *Staphylococcus aureus* strain ATCC 25923 and *Pseudomonas aeruginosa* ATCC PAO1 were overnight cultured to log phase OD_600_ = 0.3-0.6. Dilute the overnight culture 1:100 into fresh medium for biofilm assays. Add 100 µL dilutions to each well and culture it at room temperature without shaking. After 48 hr incubation, remove the media and rinse each well with 150 µL water for three times. Prepare elevated concentration of AMPs and treat the biofilm using total 150 µL volume in each well. Incubate it for 3 hr at room temperature and remove the supernatant. Rinse each well for three times using water. Add 150 µL of a 1% crystal violet in water to each well and incubate the plate at room temperature for 15 min. Rinse the plate three times with water to remove the free crystal violet. Turn the plate upside down and dry for overnight. Add 150 µL of 30% acetic acid in water to each well of the plate to solubilize the crystal violet on the cells. Incubate the plate at room temperature for 15 min. Transfer 100 µL of the solubilized crystal violet to another plate and quantify the absorbance at 550 nm using Cytation™ 5 Cell Imaging Multi-Mode Reader from BioTek.

### Drug resistant *Escherichia coli*

*Escherichia coli* strain ATCC 25922 was overnight cultured to log phase OD_600_ = 0.3-0.6. Initial bacterial cell density was prepared with ∼3 x 10^5^ CFU/mL in LB broth in 96-well plate. Bacteria were added to serially diluted antibiotics (e.g. ceftazidime, ciprofloxacin, streptomycin, and gentamicin) and co-incubated at 37 °C. After 12 hr incubation, the optical density of each well was recorded on a plate reader to determine whether they were sterilized or were at stationary phase growth. The *E. coli* which survived at the highest antibiotic concentration (below or near the MIC) was collected and cultured for another generation. This cycle is repeated for 10 times until the *e. coli* have resistant (2-fold higher MIC than its wildtype) against the antibiotics.

### Hemolysis assay

Fresh human red blood cells were obtained from Interstate Blood Bank, Inc., and thoroughly washed in PBS until the supernatant was clear. hRBC concentration was determined using a standard hemocytometer.In hemolysis assays serial dilutions of peptide were prepared, followed by the addition of 2 × 10^8^ hRBC/mL. After incubation for 1 hr at 37 °C the cells were centrifuged, and the released hemoglobin was measured by optical absorbance of the heme group (410 nm). Negative control was buffer only (0% lysis), and the positive controls were 20 µM melittin and distilled water (100% lysis). The measurements were made in triplicate.

### Molecular dynamics simulations and analysis

Unbiased all-atom MD simulations were performed and analyzed using GROMACS 5.0.4^106^ and Hippo BETA simulation packages <http://www.biowerkzeug.com, and VMD molecular visualization program^107^<http://www.ks.uiuc.edu/Research/vmd/>. The pdb structure of extended peptides (GL_5_KL_6_G, LDKL, and LDKA) were generated using Hippo BETA (see Table S1, Table S2, and Table S3). These initial structures were relaxed via 200 Monte Carlo steps, with water treated implicitly using a Generalized Born solvent.

After relaxation, the peptides were placed in all atom peptide/lipid/water systems containing model membranes with 100 mM K and Cl ions using CHARMM-GUI^108^<http://www.charmm-gui.org/>. Four helical peptides were initially placed on both interfaces of the bilayer and equilibrated and relaxed for ∼600 ns. After equilibration, the system was multiplied by 2×2 matrix in both the x and y directions and results in a bigger system with total 16 surface-bound peptides on the bilayer. The simulations were performed at 120 °C to speed up the kinetics, and we confirmed their simulated helicity with the liquid-state circular dichroism spectroscopy (**Table S2 and Figure S2**). MD simulations were performed with GROMACS 5.0.4 using the CHARMM36 force field^109^, in conjunction with the TIP3P water model^110^. Electrostatic interactions were computed using PME, and a cut-off of 10 Å was used for van der Waals interactions. Bonds involving hydrogen atoms were constrained using LINCS. The integration time-step was 2 fs and neighbor lists were updated every 5 steps. All simulations were performed in the NPT ensemble, without any restraints or biasing potentials. Water and the protein were each coupled separately to a heat bath with a time constant τ_T_ = 0.5 ps using velocity rescale temperature coupling. The atmospheric pressure of 1 bar was maintained using weak semi-isotropic pressure coupling with compressibility κ_z_ = κ_xy_ = 4.6 · 10^−5^ bar^−1^ and time constant τ_P_ = 1 ps.

### Oligomer population analysis

In order to reveal the most populated pore assemblies during the simulations, a complete list of all oligomers was constructed for each trajectory frame. An oligomer of order *n* was considered any set of *n* peptides that are in mutual contact, defined as a heavy-atom (N, C, O) minimum distance of <3.5 Å. Frequently, this definition overcounts the oligomeric state due to numerous transient surface bound (S-state) peptides that are only loosely attached to the transmembrane inserted peptides that make up the core of the oligomer. These S-state peptides frequently change position or drift on and off the stable part of the pore. To focus the analysis on true longer-lived TM pores, a cut-off criterion of 65° was introduced for the tilt angle τ of the peptides. Any peptide with τ ≥65° was considered in the S-state and removed from the oligomeric analysis. This strategy greatly reduced the noise in the oligomeric clustering algorithm by focusing on the true longer-lived pore structures. Population plots of the occupation percentage of oligomer *n* multiplied by its number of peptides *n*, were then constructed. These reveal how much peptide mass was concentrated in which oligomeric state during the simulation time.

### Membrane partitioning and secondary structure

Peptide solutions (50 µM peptide concentration) were titrated with POPC and POPG LUVs (between 0-5 mM) and the corresponding changes of tryptophan fluorescent spectra were collected (**Figure S1**). 7F3 and 28H6 show maximum fluorescent emission of ∼331 nm in phosphate buffer, suggesting aggregate formation. Other peptides have tryptophan fluorescence peaks at ∼348 nm, indicative of monomeric peptides or low multimeric soluble aggregates. Change of the maximum wavelength indicates the partitioning between water and lipid phases. It gives a direct measure of the binding free energy (*∆G*_*binding*_) for each peptide with different lipids. Binding free energy of toxic peptides (**Figure S1a-j**; 4H9, 7F3, 28H6, and 25B2), which porate both POPC and POPG LUVs, are between −5.5 and −9.5 kcal/mol. The membrane-selective peptides (**Figure S1k-r**; 7G6, 7D12, 11D12, and 24F1) have lower binding free energy toward POPC LUV (*∆G*_*binding*_ = −4.0 to −5.7 kcal/mol) than POPG LUV (*∆G*_*binding*_ = −5.7 to −7.1 kcal/mol). It shows that the strength of peptide binding is essential for membrane-selectivity and *∆G*_*binding*_ = −5.7 kcal/mol is the cut-off.

We further performed CD spectroscopy to study the secondary structure of these peptides with each POPC and POPG LUVs at elevated temperature (**Figure S2 and Table S2**). CD spectroscopy shows that all the toxic peptides are helical structure (54-75% helicity) in the solution, and membrane-selective peptides are mostly coiled structure (22-38% helicity). The secondary structure of the peptides in solution explain why these LDKA analogues have selectivity toward different membrane types and result in different binding free energy. Coiled structure exposes its intramolecular hydrogen bonds to water that make the compound more polar; in opposite, helical structure makes it more hydrophobic. Therefore, toxic peptides have higher helical content and strong interaction with both membrane types. Interestingly, 28H6 only folds beta-strand structure in POPG LUV, and the temperature at 95 °C can break the intermolecular hydrogen bonds and reverse it to helix. As expected, the membrane-selective peptides only fold helix in POPG LUV and have no response to POPC LUV. Most of the helical structures are highly resistant to thermal denaturation (at 95 °C) when they once fold in the membrane (**Figure S2**).

The linear regression analysis shows strong correlation between hydrophobic moments, helicity in POPC LUV, and ANTS/DPX leakage fraction from POPC LUV (**Figure S3a**). It confirms the interaction between peptide and POPC LUV is strongly dependent on the peptide’s hydrophobic moment; however, it does not correlate to the membrane pore size. **Figure S3b** shows that the helicity of a peptide is linearly correlated to the hydrophobic moment, which is promoted by the hydrophobicity. We further analyzed the AMPs from *APD* that have peptide length between 5 and 30 amino acids, which dominate >50% peptide population (1,500 AMPs) in *APD*. We grouped the AMPs by their peptide length and averaged each of their net charge and hydrophobic moment. It shows that increased hydrophobic moment corresponds to higher net charge (**Figure S4a**).

We analyzed the sequence of LDKA analogues and compared them to the AMPs from *APD* that have same peptide length to LDKA (**Figure S4b**). It showed that the toxic LDKA peptides have higher hydrophobic moment 3.41-4.78 than membrane-selective LDKA peptides with hydrophobic moment 1.92-3.32 (Table 1), which correspond to their specificity toward different membrane types (**Figure S4c**) and toxicity to human red blood cell (**Figure S4d**). We found hydrophobic moment 3.37 is a cut-off between membrane-selective and toxic peptides in the LDKA library peptides. However, the cut-off may shift in different peptide length and charge distribution (**Figure S4e-h and Table S5**); therefore, bigger sample size is necessary to improve this sequence-based prediction of membrane selectivity.

## Data availability

The data that support the findings of this study are available from the corresponding author on reasonable request.

## Acknowledgements

We thank the Karlsruhe Institute of Technology (KIT) ANKA synchrotron CD beamline staff for support and beamtime. We thank Jochen Bürck at KIT for valuable discussion and technical support for ANKA synchrotron CD beamline. We thank Guangshun Wang at the University of Nebraska Medical Center for providing the raw data of the antimicrobial peptide database. We thank Katherine Tripp at the Center for Molecular Biophysics, Johns Hopkins University for helping the experimental setup for isothermal titration calorimeter. We thank Jodie Franklin at the Synthesis and Sequencing Facility at the Johns Hopkins University School of Medicine for sequencing the LDKA peptides. We thank Kalina Hristova and Honggang Cui at Johns Hopkins University and Gregory Wiedman at Seton Hall University and Bonnie Wallace at Birkbeck, University of London and Matthew Upton at University of Plymouth for valuable discussions. Simulation resources were supported by the MARCC supercomputer facility at Johns Hopkins University.

## Author contributions

CHC, JPU, MBU and WCW designed the research. CHC performed most of the experiments and MD simulations. CGS performed hemolysis assay. CGS and CHC performed *in-vitro* bacterial minimum inhibitory concentration assay and bacterial biofilm assay. SG and CHC performed the minimum inhibitory concentration assay with drug resistant *E. coli*. JPU, CHC, and MBU analyzed the simulations. CHC, JPU, MBU and WCW wrote the paper, with input from the other authors.

## Additional Information

### Supplementary Information

The Supporting Information is available free of charge on the ACS Publications website.

